# Deep sequencing of primary human lung epithelial cells challenged with H5N1 influenza virus reveals a proviral role for CEACAM1

**DOI:** 10.1101/324723

**Authors:** Siying Ye, Chris Cowled, Cheng-Hon Yap, John Stambas

## Abstract

Current prophylactic and therapeutic strategies targeting human influenza viruses include vaccines and antivirals. Given variable rates of vaccine efficacy and antiviral resistance, alternative strategies are urgently required to improve disease outcomes. Here we describe the use of HiSeq deep sequencing to analyze host gene expression in primary human alveolar epithelial type II cells infected with highly pathogenic avian influenza H5N1 virus. At 24 hours post-infection, 623 host genes were significantly up-regulated, including the cell adhesion molecule *CEACAM1*. The up-regulation of *CEACAM1* was blocked in the presence of the reactive oxygen species inhibitor, apocynin. H5N1 virus infection stimulated significantly higher CEACAM1 protein expression when compared to low pathogenic PR8 H1N1 virus, suggesting a key role for CEACAM1 in influenza virus pathogenicity. Furthermore, silencing of endogenous *CEACAM1* resulted in reduced levels of proinflammatory cytokine/chemokine production, as well as reduced levels of virus replication following H5N1 infection. Our study provides evidence for the involvement of CEACAM1 in a clinically relevant model of H5N1 infection and may assist in the development of host-oriented antiviral strategies.

## IMPORTANCE

Highly pathogenic avian H5N1 influenza virus continues to pose a potential pandemic threat. Therapeutic strategies that enhance host immune responses to promote clearance of influenza virus infection are urgently needed. *CEACAM1* was significantly up-regulated in primary alveolar epithelial type II cells infected with H5N1 virus. It was one of 623 human genes identified using HiSeq deep sequencing analysis that were up-regulated following H5N1 infection. Although CEACAM1 is known for its multifaceted immune regulatory role, little is known about its contribution to host immunity. The results presented here show that CEACAM1regulates the innate immune response in a clinically relevant *in vitro* human model of H5N1 infection.

## INTRODUCTION

Influenza viruses cause acute and highly contagious seasonal respiratory disease in all age groups. Between 3-5 million cases of severe influenza-related illness and over 250 000 deaths are reported every year. In addition to constant seasonal outbreaks, highly pathogenic avian influenza (HPAI) strains, such as H5N1, remain an ongoing pandemic threat with recent WHO figures showing 454 confirmed laboratory infections and a mortality rate of 53%. It is important to note that humans have very little pre-existing immunity towards avian influenza virus strains. Moreover, there is no commercially available human H5N1 vaccine. Given the potential for H5N1 viruses to trigger a pandemic (1, 2), there is an urgent need to develop novel therapeutic interventions to combat known deficiencies in our ability to control outbreaks. Current seasonal influenza virus prophylactic and therapeutic strategies involve the use of vaccination and antivirals. Vaccine efficacy is highly variable as evidenced by a particularly severe 2017/18 epidemic and frequent re-formulation of the vaccine is required to combat ongoing mutations in the influenza virus genome. In addition, antiviral resistance has been reported for many circulating strains, including the avian influenza H7N9 virus that emerged in 2013 (3, 4). Influenza A viruses have also been shown to target and hijack multiple host cellular pathways to promote survival and replication (5, 6). As such, there is increasing evidence to suggest that targeting host pathways will influence virus replication, inflammation, immunity and pathology (6, 7). Alternative intervention strategies based on modulation of the host response could be used to supplement the current prophylactic and therapeutic protocols.

While the impact of influenza virus infection has been relatively well studied in animal models (8, 9), the human cellular responses are poorly defined due to the lack of available human autopsy material, especially from HPAI virus-infected patients. In the present study, we characterized influenza virus infection of primary human alveolar epithelial type II (ATII) cells isolated from normal human lung tissue donated by patients undergoing lung resection. ATII cells are a physiologically relevant infection model as they are a major target for influenza A viruses when entering the respiratory tract (10). Human host gene expression following HPAI H5N1 virus (A/Chicken/Vietnam/0008/04) infection of primary ATII cells was analyzed using Illumina HiSeq deep sequencing. In order to gain a better understanding of the mechanisms underlying modulation of host immunity in an antiinflammatory environment, we also analyzed changes in gene expression following HPAI H5N1 infection in the presence of the reactive oxygen species inhibitor (ROS), apocynin, a compound known to interfere with NADPH oxidase subunit assembly (5, 6).

Our HiSeq analysis described herein focused on differentially regulated genes following H5N1 infection and identified carcinoembryonic-antigen (CEA)-related cell adhesion molecule 1 (CEACAM1) as a key gene of interest. CEACAM1 (also known as BGP or CD66) is expressed on epithelial and endothelial cells (11), as well as B cells, T cells, neutrophils, NK cells, macrophages and dendritic cells (DCs) (12–14). Human CEACAM1 has been shown to act as a receptor for several human bacterial and fungal pathogens, including *Haemophilus influenza, Escherichia coli, Salmonella typhi and Candida albicans*, but has not as yet been implicated in virus entry (15–17). There is however emerging evidence to suggest that CEACAM1 is involved in host immunity as enhanced lymphocyte expression was detected in pregnant women infected with cytomegalovirus (18) and in cervical tissue isolated from patients with papillomavirus infection (19).

Eleven *CEACAM1* splice variants have been reported in humans (20). CEACAM1 isoforms (Uniprot P13688-1 to −11) can differ in the number of immunoglobulin-like domains present, in the presence or absence of a transmembrane domain and/or the length of their cytoplasmic tail (i.e. L, long or S, short). The full-length human CEACAM1 protein (CEACAM1-<u>4</u>L) consists of four extracellular domains (one extracellular immunoglobulin variable-region-like (IgV-like) domain and three immunoglobulin constant region 2-like (IgC2-like) domains), a transmembrane domain, and a long (L) cytoplasmic tail. The long cytoplasmic tail contains two immunoreceptor tyrosine-based inhibitory motifs (ITIMs) that are absent in the short form (20). The most common isoforms expressed by human immune cells are CEACAM1-4L and CEACAM1-3L (21). CEACAM1 interacts homophilically with itself (22) or heterophilically with CEACAM5 (a related CEACAM family member) (23). The dimeric state allows recruitment of signaling molecules such as SRC-family kinases, including the tyrosine phosphatase SRC homology 2 (SH2)-domain containing protein tyrosine phosphatase 1 (SHP1) and SHP2 members to phosphorylate ITIMs (24). As such, the presence or absence of ITIMs in CEACAM1 isoforms influences signaling properties and downstream cellular function. CEACAM1 homophilic or heterophilic interactions and ITIM phosphorylation are critical for many biological processes, including regulation of lymphocyte function, immunosurveillance, cell growth and differentiation (25, 26) and neutrophil activation and adhesion to target cells during inflammatory responses (27). It should be noted that CEACAM1 expression has been modulated *in vivo* using an anti-CEACAM1 antibody (MRG1) to inhibit CEACAM1-positive melanoma xenograft growth in SCID/NOD mice (28). MRG1 blocked CEACAM1 homophilic interactions that inhibit T cell effector function, enhancing the killing of CEACAM1+ melanoma cells by T cells (28). This highlights a potential intervention pathway that can be exploited in other disease processes, including virus infection.

Our results show that CEACAM1 mRNA and protein expression levels were highly elevated following HPAI H5N1 infection. Furthermore, small interfering RNA (siRNA)-mediated inhibition of CEACAM1 reduced inflammatory cytokine and chemokine production and more importantly, inhibited H5N1 virus replication in primary ATII, as well as in the continuous human type II respiratory epithelial A549 cell line. Taken together, these observations suggest CEACAM1 is an attractive candidate for modulating influenza-specific immunity. In summary, our study has identified a novel target that may influence HPAI H5N1 immunity and serves to highlight the importance of manipulating host responses as a way of improving disease outcomes in the context of virus infection.

## RESULTS

Three experimental groups were included in the HiSeq analysis of H5N1 infection in the presence or absence of the ROS inhibitor, apocynin: (i) uninfected cells treated with 1% DMSO (vehicle control) (ND), (ii) H5N1-infected cells treated with 1% DMSO (HD) and (iii) H5N1-infected cells treated with 1mM apocynin dissolved in DMSO (HA). These three groups were assessed in pairwise comparisons: ND *vs*. HD, ND *vs*. HA, and HD *vs*. HA.

### H5N1 infection and apocynin treatment induce differential expression of host genes

A total of 13,649 genes were identified with FPKM (fragments per kilobase of exon per million fragments mapped) > 1 in at least one of the three experimental groups. A total of 613 genes were significantly up-regulated and 239 genes were significantly down-regulated (*q* value < 0.05, ≥ 2-fold change) following H5N1 infection (ND *vs*. HD) (Fig. 1B; Table S1). HPAI H5N1 infection of ATII cells activated an antiviral state as evidenced by the up-regulation of numerous interferon-induced genes, genes associated with pathogen defense, cell proliferation, apoptosis, and metabolism (Table 1; Table S2). In the apocynin alone-treated (HA) group, a large number of genes were also significantly up-regulated (509 genes) or down-regulated (782 genes) (Fig. 1A; Table S1) relative to the control group. Whilst a subset of genes was differentially expressed in both the HD and HA groups, a majority of genes did not in fact overlap (Fig. 1B and C). This suggests that apocynin treatment alone in the absence of H5N1 infection can affect gene expression. Gene Ontology (GO) enrichment analysis of genes up-regulated by apocynin showed the involvement of the type I interferon signaling pathway (GO:0060337), the defense response to virus (GO:0009615), negative regulation of viral processes (GO:48525) and the response to stress (GO:0006950) (Table S2, “ND *vs* HA Up”, Up, up-regulation). Genes down-regulated by apocynin include those that are involved in cell adhesion (GO:0007155), regulation of cell migration (GO: 0030334), regulation of cell proliferation (GO:0042127), signal transduction (GO: 0007165) and oxidation-reduction processes (GO:0055114) (Table S2, “ND *vs* HA Down”; Down, down-regulation).

**FIG 1.**
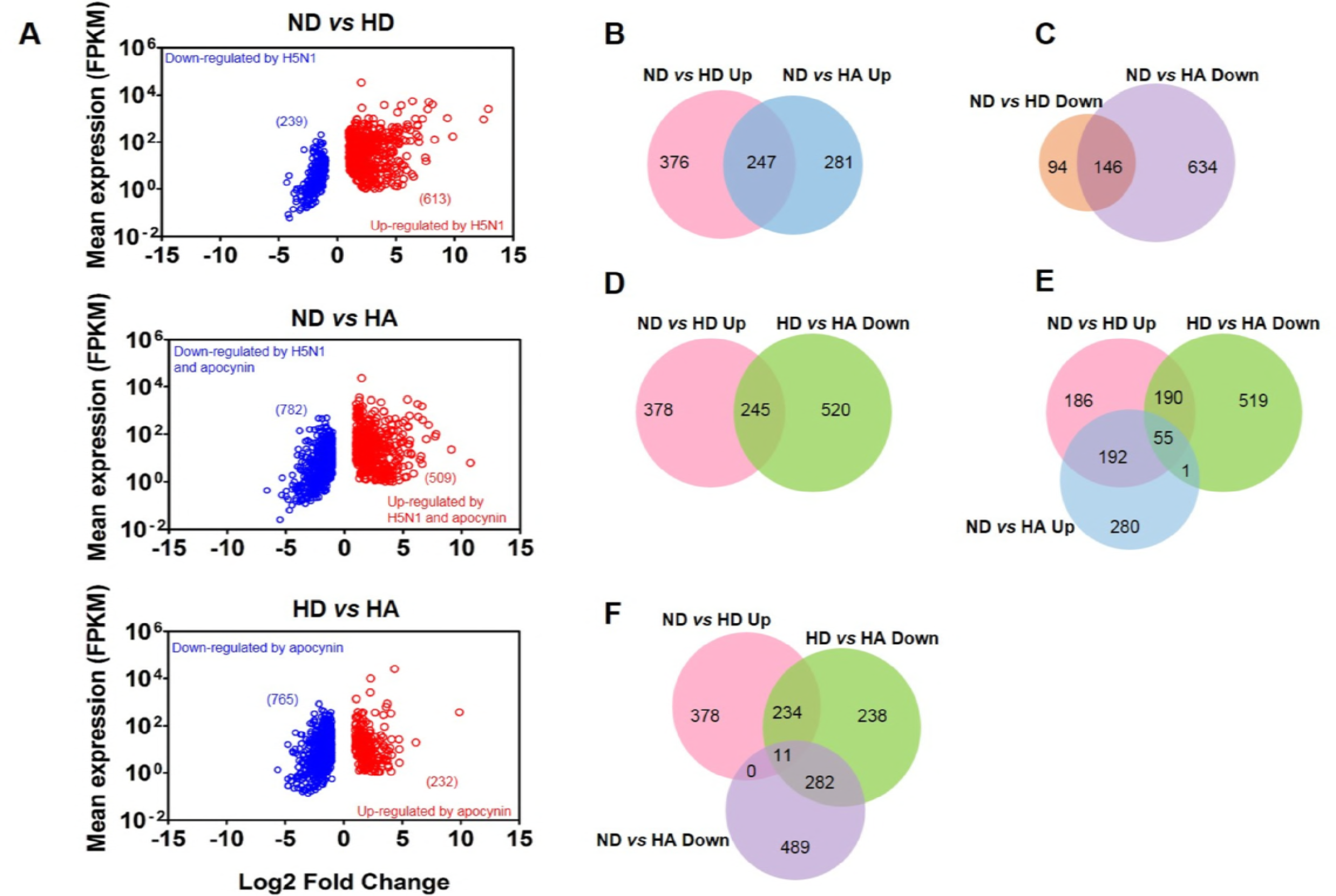
Human host gene expression profiles following HPAI H5N1 infection and apocynin treatment. (A) Genes that were statistically up-regulated (shown as red circles) or down-regulated (shown as blue circles) were assessed in pair-wise comparisons as indicated. Data are plotted as the mean FPKM of each gene obtained from HD or HA against a Log2 fold change compared to ND (HD vs. ND, HA vs. ND) or HD (HA vs. HD). (B - F) The number and overlap of genes in each pair-wise comparison is illustrated by Venn diagrams. ND, uninfected cells treated with 1% DMSO vehicle control; HD and HA, cells infected with H5N1 at a MOI of 2 for 24 hours in the presence of 1% DMSO or 1 mM apocynin, respectively. Up; up-regulated. Down; down-regulated. FPKM; fragments per kilobase of exon per million fragments mapped. The full list of transcripts identified in the HiSeq analysis, as well as differentially regulated transcripts in the three experimental groups is presented in Table S1.

**TABLE 1.** GO terms enriched in statistically significantly up-regulated genes in response to H5N1 infection (“HD *vs*. ND”) in ATII cells. The full list of GO term enrichment is presented in Table S2.

| GO Terms (Biological Processes) |  | Gene<br>Count | FDR q value |
| --- | --- | --- | --- |
| GO:0006952 | defense response | 99 | 5.00E-29 |
| GO:0043207 | response to external biotic stimulus | 87 | 1.34E-27 |
| GO:0009615 | response to virus | 49 | 3.02E-22 |
| GO:0002376 | immune system process | 145 | 6.32E-24 |
| GO:0006955 | immune response | 76 | 4.71E-20 |
| GO:0019221 | cytokine-mediated signaling pathway | 67 | 7.74E-19 |
| GO:0060337 | type I interferon signaling pathway | 22 | 1.37E-17 |
| GO:0006950 | response to stress | 177 | 1.45E-17 |
| GO:0007166 | cell surface receptor signaling pathway | 137 | 3.01E-17 |
| GO:0001817 | regulation of cytokine production | 65 | 5.99E-16 |
| GO:0045088 | regulation of innate immune response | 51 | 1.90E-15 |
| GO:0060333 | interferon-gamma-mediated signaling pathway | 20 | 3.57E-13 |
| GO:0006954 | inflammatory response | 41 | 1.36E-11 |
| GO:1903900 | regulation of viral life cycle | 25 | 4.83E-09 |
| GO:0042127 | regulation of cell proliferation | 94 | 5.01E-09 |

A total of 623 genes were up-regulated following H5N1 infection (“ND *vs* HD Up”, Fig. 1D). By overlapping the two lists of genes from “ND *vs* HD Up” and “HD *vs* HA Down”, 245 genes were shown to be down-regulated in the presence of apocynin (Fig. 1D). By overlapping three lists of genes from “ND *vs* HD Up”, “HD *vs* HA Down” and “ND *vs* HA Up”, 55 genes out of the 245 genes were present in all three lists (Fig. 1E), indicating that these 55 genes were significantly inhibited by apocynin but to a level that was still significantly higher than that in uninfected cells. The 55 genes include those involved in influenza A immunity (hsa05164; DDX58, IFIH1, IFNB1, MYD88, PML, STAT2), Jak-STAT signaling (hsa04630; IFNB1, IL15RA, IL22RA1, STAT2), cytokine-cytokine receptor interaction (hsa04060; CX3CL1, IFNB1, IL15RA, IL22RA1) and RIG-I-like receptor signaling (hsa04622; DDX58, IFIH1, IFNB1) (Table S3 and S4). Therefore, critical immune responses induced following H5N1 infection were not dampened following apocynin treatment. The remaining 190 of 245 genes excluded from the “ND vs HA Up” list were those significantly inhibited by apocynin to a level that was similar to uninfected control cells (Fig. 1E). The 190 genes that were reduced by apocynin to basal levels were those involved in PI3K-Akt signaling (hsa04151; CCND1, GNB4, IL2RG, IL6, ITGA2, JAK2, LAMA1, MYC, IPK3AP1, TLR2, VEGFC), cytokine-cytokine receptor interaction (hsa04060; VEGFC, IL6, CCL2, CXCL5, CXCL16, IL2RG, CD40, CCL5, CCL7, IL1A), TNF signaling (hsa04668; CASP10, CCL2, CCL5, CFLAR, CXCL5, END1, IL6, TRAF1, VEGFC) and apoptosis (hsa04210; BAK1, CASP10, CFLAR, CTSO, CTSS, PARP3, TRAF1) (Table S3 and S4). This is consistent with the role of apocynin in reducing inflammation and apoptosis (29).

By overlapping the three lists of genes from “ND *vs* HD Up”, “HD *vs* HA Down” and “ND *vs* HA Down”, eleven genes were found in all three comparisons. This suggests that these 11 genes are upregulated following H5N1 infection and are significantly reduced by apocynin treatment to a level lower than that observed in uninfected control cells (Fig. 1F). Among these were inflammatory cytokines/chemokines genes, including CXCL5, IL1A, AXL (a member of the TAM receptor family of receptor tyrosine kinases) and TMEM173/STING (Stimulator of IFN Genes) (Table S4).

A previous study in our laboratory demonstrated that H5N1 infection of A549 cells in the presence of apocynin enhanced expression of negative regulators of cytokine signaling, SOCS1 and SOCS3, resulting in a reduction of H5N1-stimulated cytokine and chemokine production (IL-6, IFN-β, CXCL10, and CCL5, in A549 cells) (5). We performed a qRT-PCR analysis on the same RNA samples submitted for HiSeq analysis to validate HiSeq results. *IL-6* (Fig. 2A), *IFN-β* (Fig. 2B), *CXCL10* (Fig. 2C), and *CCL5* (Fig. 2D) gene expression was significantly elevated in ATII cells following infection and was reduced by the addition of apocynin (except for *IFN-β)* (Fig. 2A-D). Consistent with previous findings in A549 cells (5), H5N1 infection alone induced the expression of *SOCS1* as shown by HiSeq and qRT-PCR analysis (Fig. 2E). Apocynin treatment further increased *SOCS1* mRNA expression (Fig. 2E). Although HiSeq analysis did not detect a statistically significant increase of *SOCS1* following apocynin treatment, the Log2 fold-changes in *SOCS1* gene expression were similar between the HD and HA groups (4.8-fold *vs* 4.0-fold) (Fig. 2E). *SOCS3* transcription was also found to be significantly increased following H5N1 infection alone following HiSeq analysis and was further increased after apocynin treatment (Fig. 2F). In contrast, *SOCS3* mRNA was only slightly increased when samples were analyzed by qRT-PCR (Fig. 2F).

**FIG 2.**
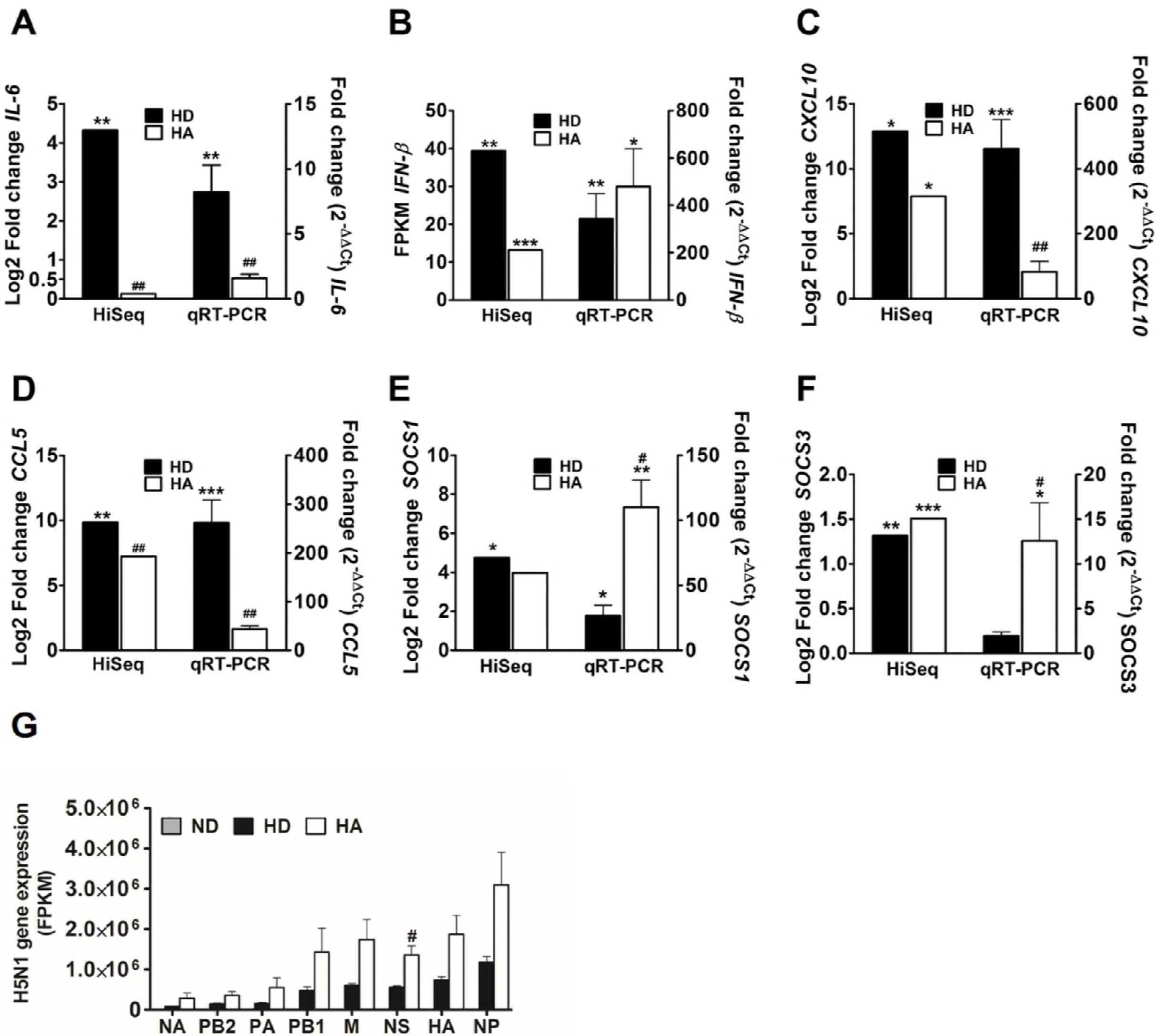
Validation of HiSeq differential gene expression using qRT-PCR. Comparison of HiSeq and qRT-PCR analysis of *IL-6* (A), *IFN-β* (B), *CXCL10* (C), *CCL5* (D), *SOCS1* (E), *SOCS3* (F) in three experimental groups of ATII cells (ND, HD and HA). Fold-changes following qRT-PCR analysis were calculated using 2^-AACt^ method (right Y axis) normalized to actin and compared with the ND group. Data from HiSeq was calculated as Log2 fold-change (left Y axis) compared with the ND group. IFN-β transcription was not detected in ND, therefore, HiSeq IFN-β data from HD and HA groups was expressed as FPKM. (G) FPKM of the eight H5N1 influenza virus genes in three experimental groups of ATII cells (ND, HD and HA). \**p* < 0.05 and \*\**p* < 0.01, \*\*\**p* < 0.001 compared with ND; ^*#*^*p* < 0.05, ^*##*^*p* < 0.01, compared with HD.

Apocynin, a compound that inhibits expression of ROS, has been shown to influence influenza-specific responses *in vitro* (5) and *in vivo* (6). Although virus titers are not affected by apocynin treatment *in vitro* (5), some anti-viral activity is observed *in vivo* when mice have been infected with a low pathogenic A/HongKong/X31 H3N2 virus (6). Our HiSeq analysis of HPAI H5N1 virus gene transcription (Fig. 2G) showed that although there was a trend for increased influenza virus gene expression following apocynin treatment, with only influenza non-structural (NS) gene expression statistically significant.

### Enrichment of antiviral and immune response genes in HPAI H5N1-infected ATII cells

GO enrichment analysis was performed on genes that were significantly upregulated following HPAI H5N1 infection in ATII cells in the presence or absence of apocynin to identify over-presented GO terms. Many of the H5N1-upregulated genes were broadly involved in defense response (GO:0006952), response to external biotic stimulus (GO:0043207), immune system processes (GO:0002376), cytokine-mediated signaling pathway (GO:0019221) and type I interferon signaling pathway (GO:0060337) (Table 1; Table S2). In addition, many of the H5N1-upregulated genes mapped to metabolic pathways (hsa01100), cytokine-cytokine receptor interaction (hsa04060), Influenza A (hsa05164), TNF signaling (hsa04668) or to Jak-STAT signaling (hsa04630) (Table S3). However, not all the H5N1-upregulated genes in these pathways were inhibited by apocynin treatment as mentioned above (Table 2; Fig. 1D; Table S3).

**TABLE 2.**
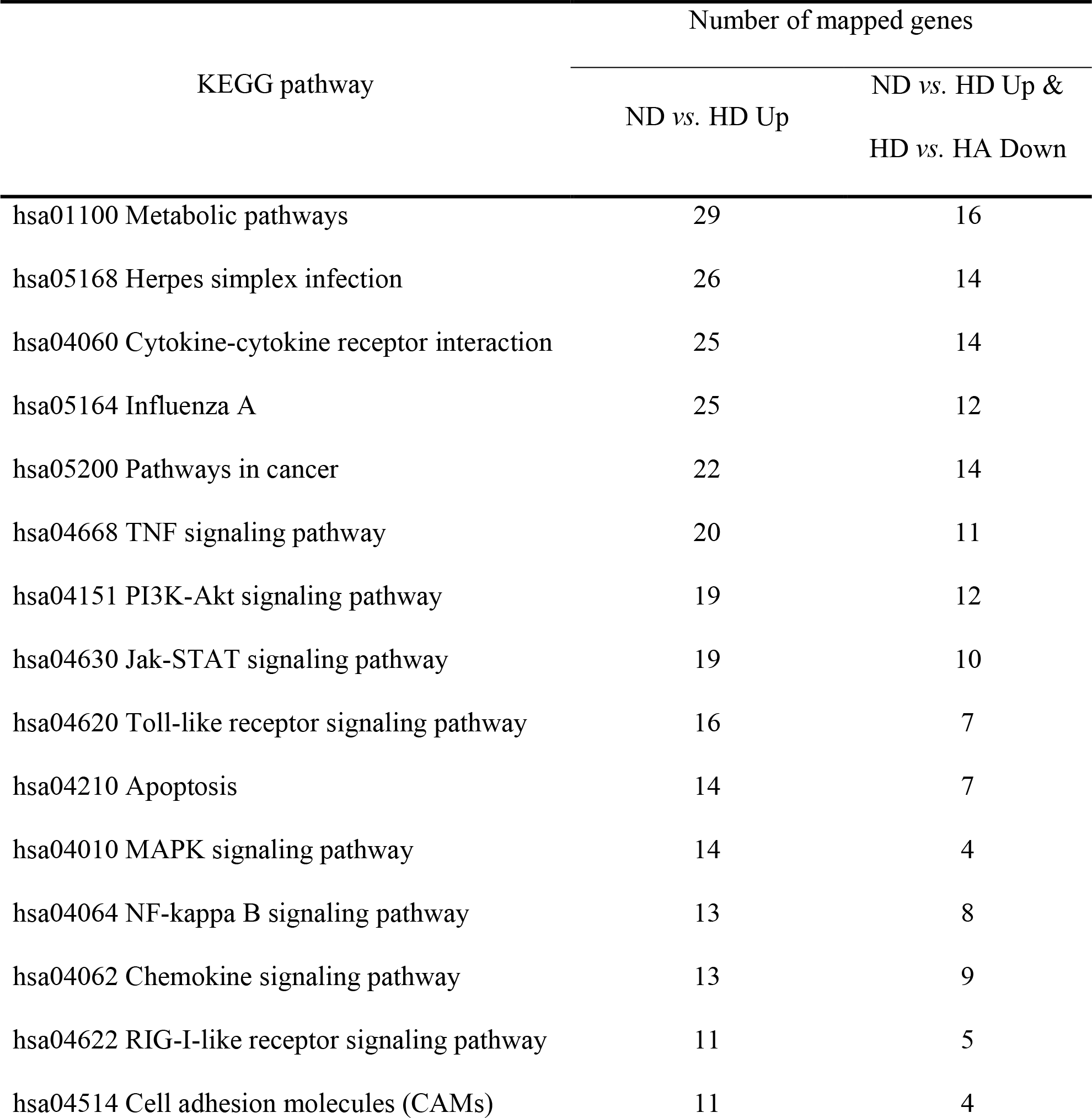
Representatives of over-represented KEGG pathways and the number of genes contributing to each pathway that is significantly up-regulated following H5N1 infection (“ND *vs*. HD Up”). The number of genes in the same pathway expressed at a significantly lower level in the HA group are also listed (Fig. 1D, “ND *vs*. HD Up and HD *vs*. HA Down”). The full list of over-represented KEGG pathways is presented in Table S3.

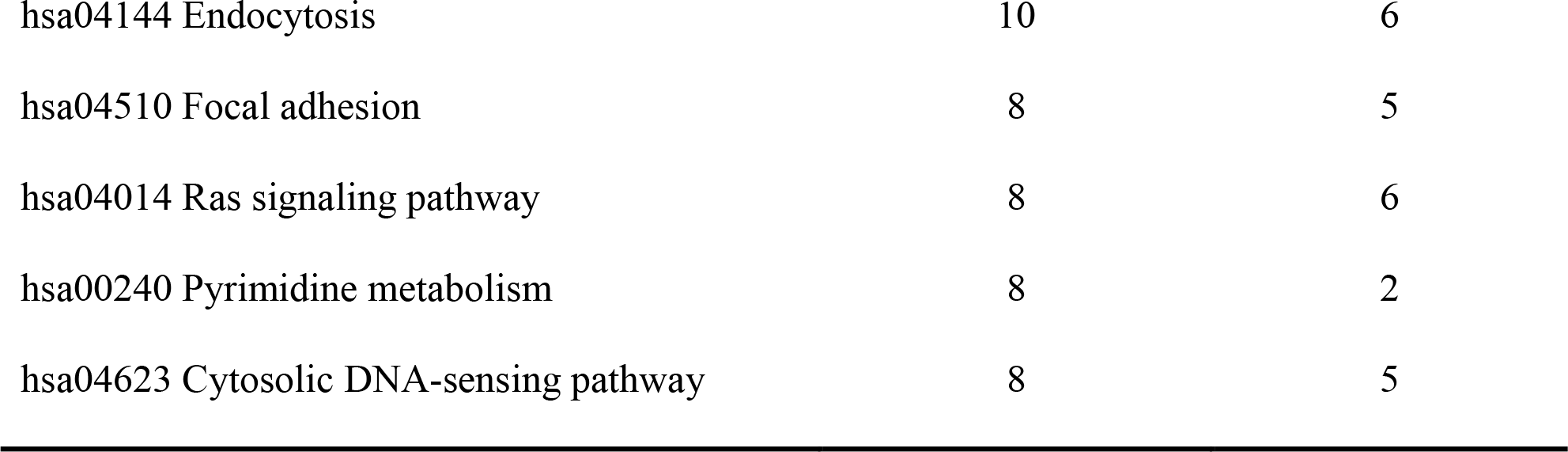

### Upregulation of the cell adhesion molecule *CEACAM1* in H5N1-infected ATII cells

The cell adhesion molecule CEACAM1 has been shown to be critical for the regulation of immune responses during infection, inflammation and cancer (20). *CEACAM1* transcript was significantly up-regulated following H5N1 infection (Fig. 3A). In contrast, a related member of the CEACAM family, *CEACAM5*, were not affected by H5N1 infection (Fig. 3B). It is also worth noting that more reads were obtained for *CEACAM5* (>1000 FPKM) than *CEACAM1* (~ 7 FPKM) in uninfected ATII cells (Fig. 3A and B), which is consistent with their normal expression patterns in human lung tissue (30). AlthoughCEACAM1 forms heterodimers with CEACAM5 (23), the higher basal expression of *CEACAM5* in ATII cells may explain why its expression was not altered by H5N1 infection. Apocynin treatment significantly reduced transcription levels of both *CEACAM1* (Fig. 3A) and *CEACAM5* (Fig. 3B). Expression of 11 *CEACAM1* variants in ATII and/or A549 cells was further characterized using qRT-PCR using primer pairs designed to specifically detect each variant (Table S5). qRT-PCR analysis confirmed up-regulation of most *CEACAM1* variants following H5N1 infection in ATII cells with *CEACAM1-4L*, *−4S* and *−3C2* increasing by > 20-fold when compared to uninfected control (Fig. 3C). mRNA transcripts of *CEACAM1-3AL* and *CEACAM1-3AS* were not expressed at sufficient levels to allow for reliable quantification by qRT-PCR. We next examined basal expression of *CEACAM1* variants. The continuous human A549 cell line is also a type II lung epithelial cell. The expression profile of *CEACAM1* variants is similar in both the A549 cell line and primary ATII cells with *CEACAM1-4L*, *−4S*, *−3L*, *−3S*, *−1L*, *−3* and *Isoform 10* readily detected (including the two common isoforms, CEACAM1-4L and −3L, in immune cells) (Fig. 3D). Endogenous CEACAM1 protein expression was also analyzed in uninfected or influenza virus-infected A549 (Fig. 3E) and ATII cells (Fig. 3F). CEACAM1 protein expression was significantly increased in A549 cells infected with PR8 H1N1 virus for 24 or 48 hours when compared to uninfected cells (Fig. 3E). Although no significant difference in CEACAM1 protein levels were observed at various MOIs (2, 5 or 10), CEACAM1 protein expression at 48 hpi was significantly higher than that at observed at 24 hpi (Fig. 3E).

**FIG 3.**
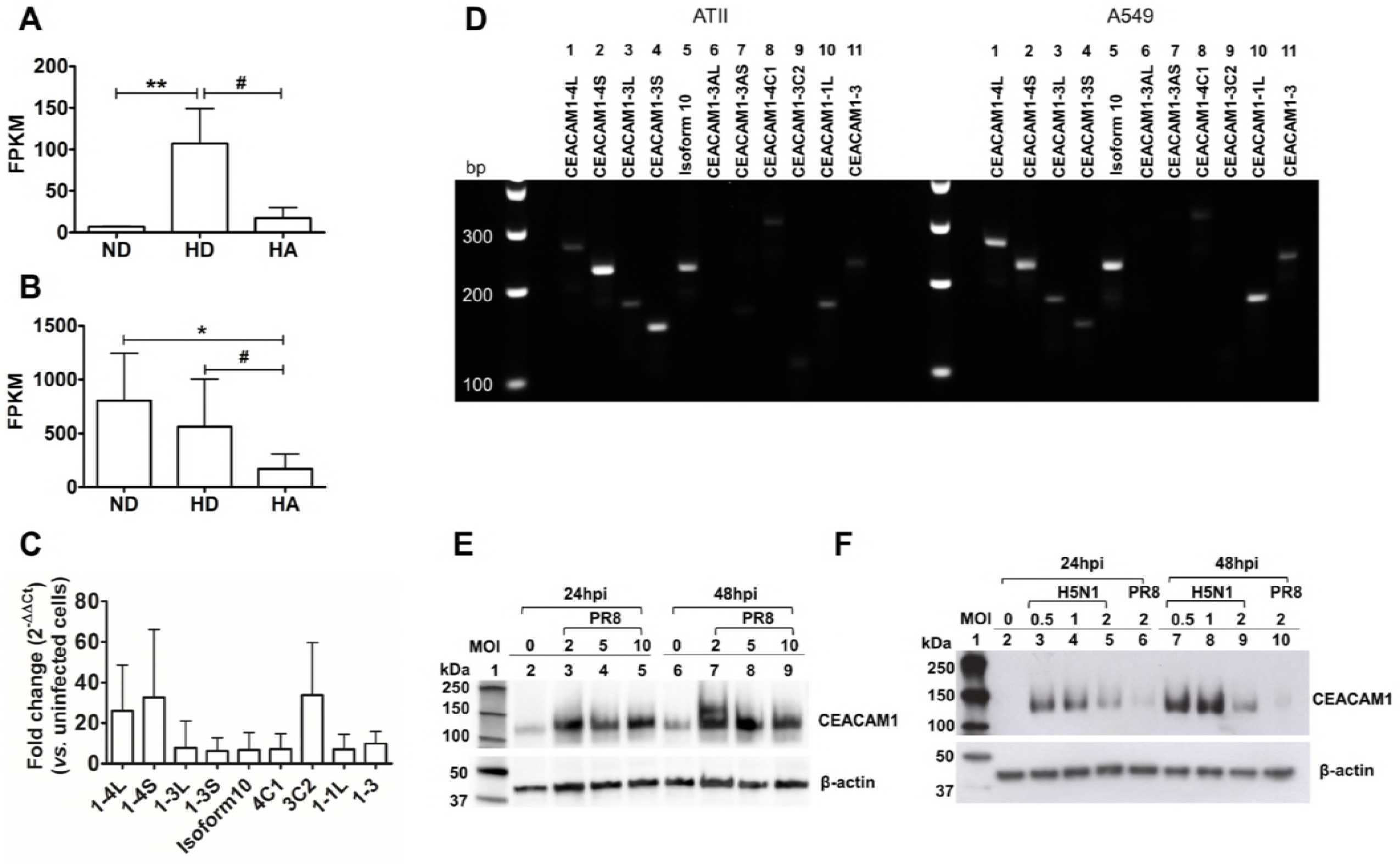
Upregulation of CEACAM1 in influenza virus-infected cells. (A) HiSeq analysis showed elevated transcription of *CEACAM1* in H5N1-infected ATII cells (HD) when compared to uninfected cells (ND). CEACAM expression was reduced in the presence of 1 mM apocynin (HA). (B) The transcription level of a related CEACAM family member, *CEACAM5*, was not altered following H5N1 infection, but was reduced by apocynin. (C) SYBR Green qRT-PCR analysis of CEACAM1 variants in ATII cells infected with H5N1 at a MOI of 2, 24 hpi. mRNAs of CEACAM1-3AL and CEACAM1-3AS were not expressed at sufficient levels to allow for reliable quantification analysis by qRT-PCR and are not shown. (D) SYBR Green qRT-PCR analysis of *CEACAM1* variants in uninfected ATII and A549 cells. qRT-PCR products (15 μL) analyzed by 2% agarose gel electrophoresis showed that both cell types expressed the same variants. *CEACAM1-3AL*, *−3AS*, *−3C2* are not readily detected. Predicted PCR product size: (1) *CEACAM1-4L*, 266 bp; (2) *CEACAM1-4S*, 245 bp; (3) *CEACAM1-3L*, 177 bp; (4) *CEACAM1-3S*, 145 bp; (5) *Isoform 10*, 233bp; (6) *CEACAM1-3AL*, 274bp; (7) *CEACAM1-3AS*, 237bp; (8) *CEACAM1-4C1*, 305bp; (9) *CEACAM1-3C2*, 109bp; (10) *CEACAM1-1L*, 173bp; (11) *CEACAM1-3*, 235 bp. (E) Endogenous CEACAM1 protein expression in A549 cells infected with PR8 virus was a maximum of 4-fold higher than uninfected cells at MOIs of 2 and 5, 48 hpi . (F) Endogenous CEACAM1 protein expression in primary human ATII cells infected with PR8 or HPAI H5N1 virus. \**p* < 0.05, \*\**p* < 0.01, compared with ND; ^*#*^*p* < 0.05, compared with HD.

After confirming the upregulation of CEACAM1 protein expression following infection with the low pathogenic PR8 virus in A549 cells, CEACAM1 protein expression was then examined in primary human ATII cells infected with HPAI H5N1 and compared to PR8 virus infection (Fig. 3F). Lower MOIs of 0.5, 1 and 2 HPAI H5N1 were tested due to the strong cytopathogenic effect it causes at higher MOIs. Endogenous CEACAM1 protein levels were significantly elevated in H5N1-infected ATII cells at all three MOIs tested. Similar levels of CEACAM1 protein expression were observed at MOIs of 0.5 and 1 and were higher at 48 hpi when compared to 24 hpi (Fig. 3F). HPAI H5N1 virus infection at MOIs of 0.5, 1 and 2 stimulated higher endogenous levels of CEACAM1 protein expression when compared to low pathogenic PR8 H1N1 virus infection at a MOI of 2 (a maximum 23-fold increase induced by H5N1 at MOIs of 0.5 and 1, 48 hpi when compared to PR8 at MOI of 2), suggesting a possible role for CEACAM1 in influenza virus pathogenicity (Fig. 3F).

### Knockdown of endogenous CEACAM1 inhibits H5N1 replication

In order to understand the role of CEACAM1 in influenza pathogenesis, A549 and ATII cells were transfected with siCEACAM1 to knockdown endogenous CEACAM1 protein expression. ATII and A549 cells were transfected with siCEACAM1 or siNeg negative control and expression of *CEACAM1* variants and CEACAM1 protein were analyzed using SYBR Green qRT-PCR and Western blotting. SYBR Green qRT-PCR analysis showed that ATII cells transfected with 15 pmol of siCEACAM1 significantly reduced expression of *CEACAM1-4L*, *−4S* and *Isoform 10* when compared to siNeg control (Fig. 4A). Transfection with siNeg significantly increased *Isoform 10* and resulted in marginal increases in *CEACAM1-3AL*, *−4C1* and *−1L* when compared to mock transfection controls. *CEACAM1-3L* was also slightly increased following siCEACAM1 transfection (Fig. 4A; Fig. 4B, white arrows). Equal volume of SYBR Green qRT-PCR amplification reaction (without normalization against the house-keeping gene, β-actin) were also subjected to agarose gel electrophoresis for visualization and showed specific amplification of each variant and complete knockdown of *CEACAM1-4L* and reduced expression of *CEACAM1-4S* and *Isoform 10* (Fig. 4B, black arrows). Similar results were also observed in A549 cells (data not shown). CEACAM1 protein expression was reduced by approximately 50% in both ATII and A549 cells following siCEACAM1 transfection when compared with siNeg-transfected cells (Fig. 4C).

**FIG 4.**
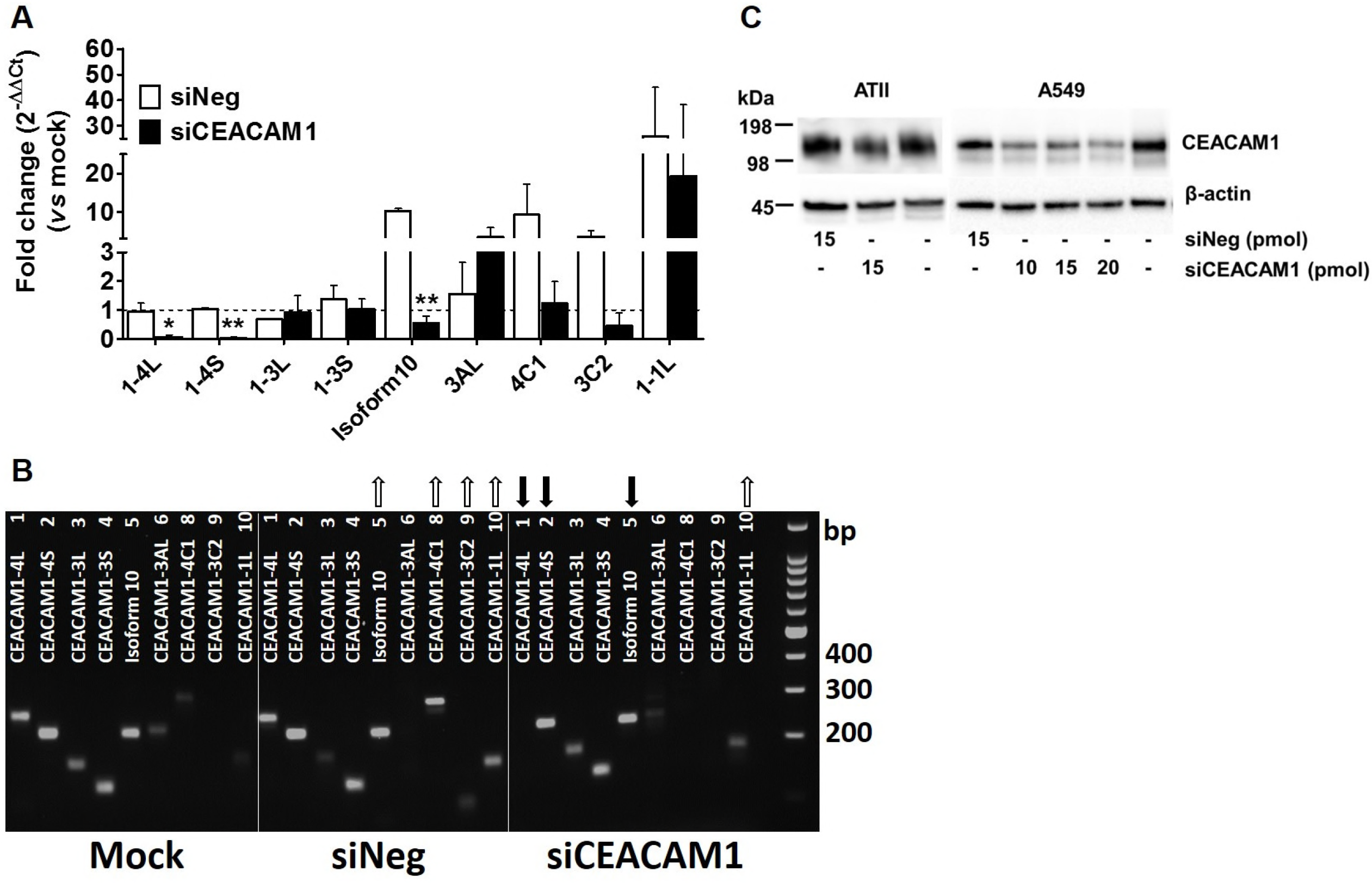
siRNA-mediated silencing of *CEACAM1* in ATII and A549 cells. (A) SYBR Green qRT-PCR analysis of siRNA-mediated silencing of endogenous *CEACAM1* in ATII cells. Transfection with 15 pmol of siCEACAM1 resulted in significant down-regulation of *CEACAM1-4L* (#1), *−4S* (#2) and *Isoform 10* (#5) compared to transfected with siNeg controls. Data are expressed as fold-change (2^−ΔΔCt^ method) normalized to actin and compared with mock-transfected cells. Dotted line indicates 1. \**p* < 0.05, \*\**p* < 0.01. (B) Equal volume of qRT-PCR reactions (15 μL) were analyzed and visualized on 2% agarose gel electrophoresis. Down-regulated *CEACAM1-4L* (#1), *−4S* (#2) and *Isoform 10* (#5) in siCEACAM1-transfected cells is shown with black arrows. When compared to mock-transfected cells, expression of *Isoform 10* (#5), *−4C1* (#8), *−3C2* (#9) and *−1L* (#10) was slightly increased in siNeg-transfected cells and *CEACAM1-1L* (#10) was also slightly increased in siCEACAM1-transfected cells as indicated by the white arrows. *CEACAM1-3AS* (#7) and *−3* (#11) were not detected and are not shown. (C) Western blot analysis of siCEACAM1-mediated knockdown of endogenous CEACAM1 in ATII and A549 cells transfected with siCEACAM1 in comparison to mock-transfected cells and cells transfected with siNeg control.

It is important to note that the anti-CEACAM1 antibody only detects L isoforms based on epitope information provided by Abcam. Therefore, observed reductions in CEACAM1 protein expression can be attributed mainly to the abolishment of *CEACAM1-4L* isoform. Increasing doses of siCEACAM1 (10, 15 and 20 pmol) did not further down-regulate CEACAM1 protein expression in A549 cells. As such, 15 pmol of siCEACAM1 was chosen for subsequent knockdown studies in both ATII and A549 cells (Fig. 4C).

The functional consequences of CEACAM1 knockdown were then examined in ATII and A549 cells following H5N1 infection. IL-6, IFN-β, CXCL10, CCL5, and TNF production was analyzed in H5N1-infected A549 and ATII cells using qRT-PCR. ATII (Fig. 5A) and A549 cells (Fig. 5B) transfected with siCEACAM1 showed significantly lower expression of cytokines and chemokines, including IL-6, CXCL10, CCL5 and TNF-α in A549 cells, when compared with siNeg-transfected cells. However, the expression of the anti-viral cytokine, IFN-β, was not affected in both cells types. In addition, the TNF-α expression, which can be induced type I IFNs (31), was also not affected in siCEACAM1-transfected ATII cells (Fig. 5A). Hypercytokinemia or “cytokine storm” in H5N1 and H7N9 virus-infected patients is thought to contribute to inflammatory tissue damage (32, 33). Down-regulation of CEACAM1 in the context of severe viral infection may reduce inflammation caused by H5N1 infection without dampening the antiviral response. Furthermore, virus replication was significantly reduced by 5.2-fold in ATII (Fig. 5C) and 4.8-fold in A549 cells (Fig. 5D) transfected with siCEACAM1 when compared with siNeg-transfected controls cells. Virus titers in siNeg-transfected control cells were not significantly different from those observed in mock-transfected control cells (Fig. 5C and D).

**FIG 5.**
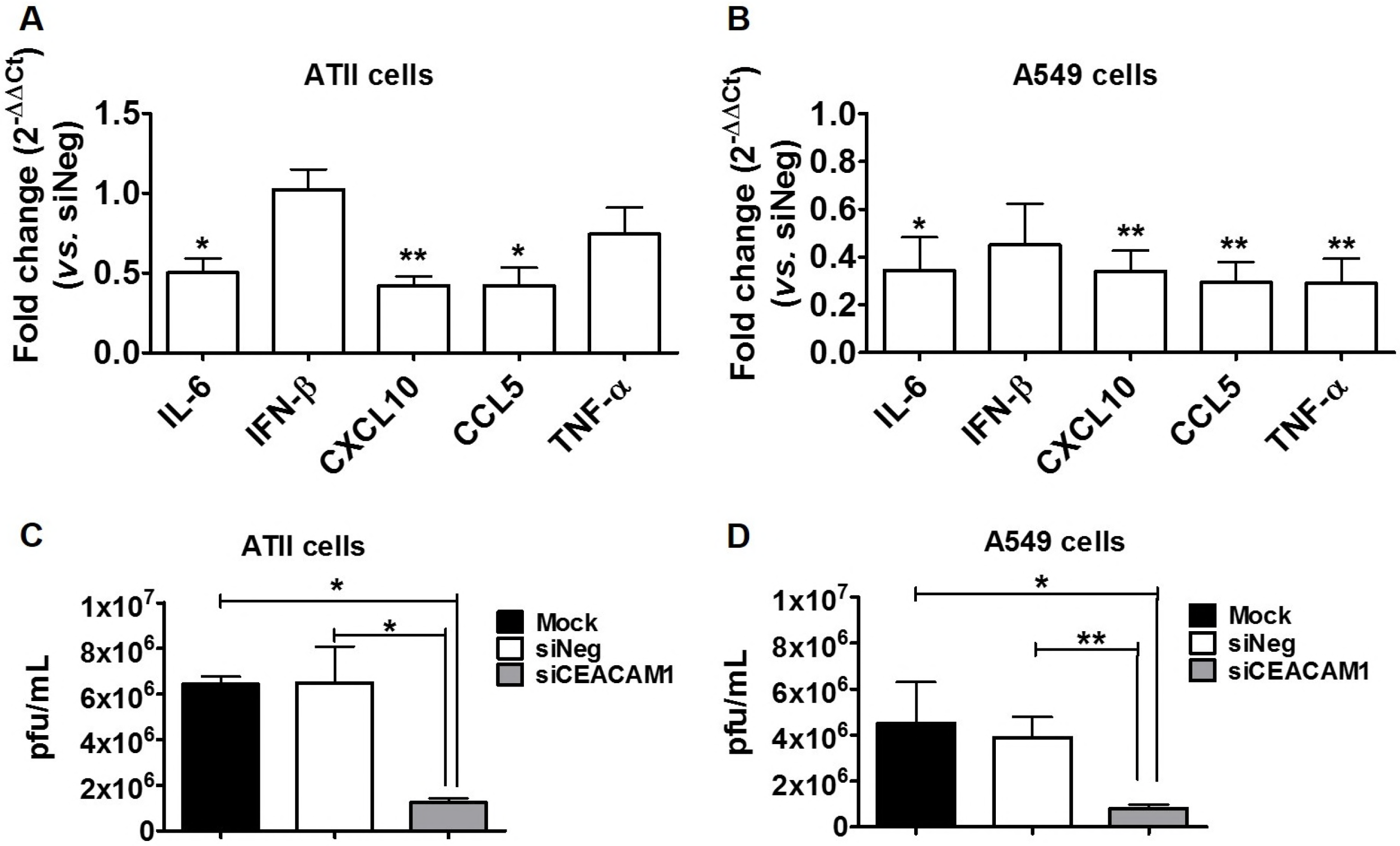
Knockdown of CEACAM1 inhibits H5N1-induced inflammation and virus replication. ATII cells (A) and A549 cells (B) transfected with 15 pmol siCEACAM1 showed reduced cytokine and chemokine production in response to H5N1 infection (MOI of 0.01, 24 hpi) by qRT-PCR com-pared to siNeg-transfected control cells. One sample t test compared to siNeg-transfected cells that have a fold-change of 1, \**p* < 0.05 and \*\**p* < 0.01. Plaque assay of H5N1 virus titres in ATII (C) and A549 (D) that were either mock-transfected or transfected with siCEACAM1 or siNeg. \**p* < 0.05 and \*\**p* < 0.01.783

### DISCUSSION

Influenza viruses utilize host cellular machinery to manipulate normal cell processes in order to promote replication and evade host immune responses. Studies in the field are increasingly focused on understanding and modifying key host factors in order to ameliorate disease. Examples include modulation of reactive oxygen species (ROS) to reduce inflammation (5) and inhibition of NFκB and mitogenic Raf/MEK/ERK kinase cascade activation to suppress viral replication (34, 35). These host targeting strategies will offer an alternative to current interventions that are focused on targeting the virus. In the present study, we analyzed human host gene expression profiles following HPAI H5N1 infection and treatment with the antioxidant apocynin. As expected, genes that were significantly upregulated following H5N1 infection were involved in biological processes including cytokine signaling, immunity and apoptosis. In addition, H5N1-upregulated genes were also involved in regulation of protein phosphorylation, cellular metabolism and cell proliferation, which are thought to be exploited by viruses for replication (36). Apocynin treatment had both anti-viral (Table S2, S3, S4) (6) and pro-viral impact (Fig. 2G), which is not surprising as ROS are potent microbicidal agents, as well as important immune signaling molecules at different concentrations (37). In our hands, apocynin treatment reduced H5N1-induced inflammation, but unavoidably impacted the cellular defense response, cytokine production and cytokine-mediated signaling. Importantly, critical antiviral responses were not compromised, i.e. the expression of pattern recognition receptors (e.g. *DDX58* (RIG-I), *TLRs*, *IFIH1* (MDA5)) was not down-regulated (Table S1). Given the significant interference of influenza viruses on host immunity, we focused our attention on key regulators of the immune responses. Through HiSeq analysis, we identified the cell adhesion molecule CEACAM1 as a critical regulator of immunity. Knockdown of endogenous CEACAM1 not only reduced H5N1-stimulated inflammatory cytokine/chemokine production but also inhibited virus replication.

H5N1 infection resulted in significant upregulation of a number of inflammatory cytokines/chemokines genes, including AXL and STING, which were significantly reduced by apocynin treatment to a level lower than that observed in uninfected cells (Table S4). It has been previously demonstrated that treatment of PR8 H1N1-infected mice with anti-AXL antibody significantly reduces lung inflammation and virus titers (38). STING has been shown to be important for promoting anti-viral responses, as STING-knockout THP-1 cells produce less type I IFN following influenza A virus infection (39). Reduction of STING gene expression or other anti-viral factors (e.g. IFNB1, MX1, ISG15; Table S1) by apocynin, may in part, explain the slight increase of the transcription levels of all influenza genes (NS gene was significantly upregulated) following apocynin treatment (Fig. 2G). These results also suggest that apocynin treatment may reduce H5N1-induced inflammation and apoptosis. Indeed, the anti-inflammatory and anti-apoptotic effects of apocynin have been shown previously in a number of disease models, including diabetes mellitus (40), myocardial infarction (41), neuroinflammation (42) and influenza virus infection (5).

Recognition of intracellular viral RNA by pattern recognition receptors (PRRs) triggers the release of pro-inflammatory cytokines/chemokines that recruit innate immune cells, such as neutrophils and NK cells to the site of infection to assist in viral clearance (43). Neutrophils exert their cytotoxic function by first attaching to influenza-infected epithelial cells via adhesion molecules, such as CEACAM1 (44). Moreover, studies have indicated that influenza virus infection promotes neutrophil apoptosis (45), delaying virus elimination (46). Phosphorylation of CEACAM1 ITIM motifs and activation of caspase-3 is critical for mediating anti-apoptotic events and for promoting survival of neutrophils (27). This suggests that CEACAM1-mediated anti-apoptotic events may be important for the resolution of influenza virus infection.

NK cells play a critical role in innate defense against influenza virus by recognizing and killing infected cells. Influenza viruses, however, employ several strategies to escape NK effector functions, including modification of influenza hemagglutinin (HA) glycosylation to avoid NK activating receptor binding (47). Homo-or heterophilic CEACAM1 interactions have been shown to inhibit NK-killing (25, 26), and are thought to contribute to tumor cell immune evasion (48). Given these findings, one could suggest the possibility that upregulation of CEACAM1 (to inhibit NK activity) may be a novel and uncharacterized immune evasion strategy employed by influenza viruses. Our laboratory is now investigating the role of CEACAM1 in NK cell function. Small-molecule inhibitors of protein kinases or protein phosphatases (e.g. inhibitors for Src, JAK, SHP2) have been developed as therapies for cancer, inflammation, immune and metabolic diseases (49). Modulation of CEACAM1 phosphorylation, dimerization and the downstream function with small-molecule inhibitors may assist in dissecting the contribution of CEACAM1 to NK cell activity.

It has also been shown that another related CEACAM family member, CEACAM6, is up-regulated following influenza infection and can interact with influenza NA protein in A549 and NHBE cells (50). Through this interaction, influenza viruses promote cell survival and viral replication by activating the Src/Akt signaling pathway. Moreover, silencing of CEACAM6 suppressed PR8 virus replication in A549 cells by inhibiting phosphorylation and activation of Src and Akt (50). Although both CEACAMs can be found in lung epithelial cells (30), unlike CEACAM1, CEACAM6 lacks an intracellular C-terminal tail (20) and differs from CEACAM1 in modulating cellular signaling events. In support of this notion, we did not observe a difference in apoptosis of PR8-infected A549 cells following transfection with siCEACAM1, siNeg, or mock-transfected cells using flow cytometry-based Annexin V/7AAD staining (data not shown). Our study, together with the CEACAM6 findings, serve to highlight the emerging role of CEACAMs in influenza virus infection.

The molecular mechanism of CEACAM1 action following infection has also been explored in A549 cells using the low pathogenic PR8 virus (51). Vitenshtein *et al*. demonstrated that CEACAM1 was up-regulated following recognition of viral RNA by RIG-I, and that this up-regulation was interferon regulatory factor 3 (IRF3)-dependent. In addition, phosphorylation of CEACAM1 by SHP2 inhibited viral replication by reducing phosphorylation of mammalian target of rapamycin (mTOR) to suppress global cellular protein production. In the present study, we used a more physiologically relevant infection model, primary human ATII cells, to study the role of CEACAM1 in influenza virus infection, focusing on HPAI H5N1 virus. Consistent with findings from Vitenshtein *et al.*, significant up-regulation of CEACAM1 protein was observed following influenza virus infection, especially in HPAI H5N1-infected cells. However, in contrast to the inhibitory effects of CEACAM1 on influenza virus replication observed by Vitenshtein *et al.*, knockdown of endogenous CEACAM1 protein expression reduced HPAI H5N1 titers by 4.8-fold in ATII cells. Despite the use of two different *in vitro* experimental settings, different influenza virus strains, infection doses and time points, both studies agree that CEACAM1 plays an important role in influenza virus infection and warrants further investigation.

Further studies will be required to investigate/confirm the molecular mechanisms of CEACAM1 up-regulation following influenza virus infection, especially *in vivo*. As up-regulation of CEACAM1 has been observed in other virus infections, such as cytomegalovirus (18) and papillomavirus (19), it will be important to determine whether a common mechanism of action can be attributed to CEACAM1 in order to determine its functional significance. If this can be established, CEACAM1 could be used as a target for the development of a pan-antiviral agent. Furthermore, it has been shown that CEACAM1-S isoforms and soluble isoforms (CEACAM1-4C1, -3, -3C2) also have regulatory effects (stimulatory or inhibitory) on CEACAM1-L isoforms (20, 52). Although CEACAM1-Ls are the dominate isoforms, the involvement of other CEACAM1 isoforms in influenza immunity should also be further investigated, especially as H5N1 infection increased mRNA expression levels of several *CEACAM1* variants (Fig. 3C).

In summary, molecules on the cell surface such as CEACAM1 are particularly attractive candidates for therapeutic development, as drugs do not need to cross the cell membrane in order to be effective. Targeting of host-encoded genes in combination with current antivirals and vaccines may be a way of reducing morbidity and mortality associated with influenza virus infection. Our study clearly demonstrates that increased CEACAM1 expression is observed in primary human ATII cells infected with highly pathogenic H5N1 influenza virus. Importantly, knockdown of CEACAM1 expression resulted in reduced influenza virus replication and suggests targeting of this molecule may assist in improving disease outcomes.

## MATERIALS AND METHODS

### Isolation and culture of primary human ATII cells

Human non-tumor lung tissue samples were donated by anonymous patients undergoing lung resection at Geelong Hospital, Barwon Health, Australia. The research protocols and human ethics were approved by the Human Ethics Committees of Deakin University, Barwon Health and the Commonwealth Scientific and Industrial Research Organisation (CSIRO). The sampling of normal lung tissue was confirmed by the Victorian Cancer Biobank, Australia. Human alveolar epithelial type II (ATII) cells were isolated and cultured using a previously described method (53, 54) with minor modifications. Briefly, lung tissue with visible bronchi was removed and perfused with abundant PBS and submerged in 0.5% Trypsin-EDTA (Gibco) twice for 15 min at 37°C. The partially digested tissue was sliced into sections and further digested in Hank’s Balanced Salt Solution (HBSS) containing elastase (12.9 units/mL; Roche Diagnostics) and DNase I (0.5 mg/mL; Roche Diagnostics) for 60 min at 37°C. Single cell suspensions were obtained by filtration through a 40 μm cell strainer and cells were allowed to attach to a petri dish in a 1:1 mixture of DMEM/F12 medium (Gibco) and small airway growth medium (SAGM) medium (Lonza) containing 5% fetal calf serum (FCS) and 0.5 mg/mL DNase I for 2 hours at 37°C. The non-adherent cells, including ATII cells, were collected and subjected to centrifugation at 300 g for 20 min on a discontinuous Percoll density gradient (1.089 and 1.040 g/mL). Purified ATII cells from the interface of two density gradients was collected, washed in HBSS, and re-suspended in SAGM media supplemented with 1% charcoal-filtered FCS (Gibco) and 100 units/mL penicillin and 100 μg/mL streptomycin (Gibco), prior to plating on polyester Transwell inserts (0.4 μm pore; Corning) coated with type IV human placenta collagen (0.05 mg/mL; Sigma) at 300,000 cells/cm^2^. Cells were then cultured under liquid-covered conditions in a humidified incubator (5% CO_2_, 37°C) and growth medium was changed every 48 hours. These culture conditions encourage ATII cells to grow and differentiate into alveolar epithelial cells.

### Cell culture and media

A549 carcinomic human alveolar basal epithelial type II cells and Madin-Darby canine kidney (MDCK) cells were provided by the tissue culture facility of CSIRO Australian Animal Health Laboratory (AAHL). A549 and MDCK cells were maintained in Ham’s F12K medium (GIBCO) and RPMI-1640 medium (Invitrogen), respectively, supplemented with 10% FCS, 100 U/mL penicillin and 100 μg/mL streptomycin (GIBCO) and maintained at 37°C, 5% CO_2_.

### Virus and viral infection

HPAI A/chicken/Vietnam/0008/2004 H5N1 (H5N1) was obtained from CSIRO Australian Animal Health Laboratory (AAHL). Viral stocks of A/Puerto Rico/8/1934 H1N1 (PR8) were obtained from the University of Melbourne. Virus stocks were prepared using standard inoculation of 10-day-old embryonated eggs. A single stock of virus was prepared for use in all assays. All H5N1 experiments were performed within biosafety level 3 laboratories (BSL3) at CSIRO AAHL.

For HiSeq analysis, ATII cells were infected on the apical side with H5N1 at a multiplicity of infection (MOI) of 2 for 24 hours as previously described (5, 55) in serum-free SAGM media supplemented with 0.3% bovine serum albumin (BSA) containing 1 mM apocynin dissolved in DMSO or 1% DMSO vehicle control. Uninfected ATII cells incubated in media containing 1% DMSO were used as a negative control. For other subsequent virus infection studies, cells were infected with influenza A viruses at various MOIs as indicated in the text. For PR8 infection studies, a final concentration of 0.5 μg/mL L-1-Tosylamide-2-phenylethyl chloromethyl ketone (TPCK)-treated trypsin (Worthington) was included in media post-inoculation to assist replication. Virus titers were determined using standard plaque assays using MDCK cells as previously described (56).

### RNA extraction, quality control (QC) and HiSeq analysis

ATII cells from three individual donors were used for HiSeq analysis. Total RNA was extracted from cells using a RNeasy Mini kit (Qiagen). Influenza-infected cells were washed with PBS three times and cells lysed with RLT buffer supplemented with β-mercaptoethanol (10 μL/mL; Gibco). Cell lysates were homogenized with QIAshredder columns followed by on-column DNA digestion with the RNase-Free DNase Set (Qiagen), and RNA extracted according to manufacturer’s instructions. Initial QC was conducted to ensure that the quantity and quality of RNA samples for HiSeq analysis met the following criteria; 1) RNA samples had OD260/280 ratios between 1.8 and 2.0 as measured with NanoDrop™ Spectrophotometer (Thermo Scientific); 2) Sample concentrations were at a minimum of 100 ng/μl; 3) RNA was analyzed by agarose gel electrophoresis. RNA integrity and quality were validated by the presence of the sharp clear bands of 28S and 18S ribosomal RNA, with a 28S:18S ratio of 2:1, along with the absence of genomic DNA and degraded RNA. As part of the initial QC and as an indication of consistent H5N1 infection, parallel quantitative real-time reverse transcriptase PCR (qRT-PCR) was performed as previously described (5) to measure mRNA expression of *IL-6*, *IFN-β*, *CXCL10*, *CCL5*, suppressor of cytokine signaling *(SOCS*) 1 and *SOCS3* (Fig. 2), which is known to be up-regulated following HPAI H5N1 infection of A549 cells (5). RNA samples were stored in 0.1 volumes of 3 M Sodium Acetate (pH7.5 in DEPC-treated water) and 2 volumes of 100% Ethanol and submitted to Macrogen Inc. (Seoul, Republic of Korea) for HiSeq analysis.

### Sequencing analysis and annotation

After confirming checksums and assessing raw data quality of the FASTQ files with FASTQC, RNA-Seq reads were processed according to standard Tuxedo pipeline protocols (57), using the annotated human genome (GRCh37, downloaded from Illumina iGenomes) as a reference. Briefly, raw reads for each sample were mapped to the human genome using TopHat2, sorted and converted to SAM format using Samtools and then assembled into transcriptomes using Cufflinks. Cuffmerge was used to combine transcript annotations from individual samples into a single reference transcriptome, and Cuffquant was used to obtain per-sample read counts. Cuffdiff was then used to conduct differential expression analysis. All programs were run using recommended parameters, however the reference gtf file provided to cuffmerge was first edited using a custom python script to exclude lines containing features other than exon/cds, and contigs other than chromosomes 1-22, X, Y.

### Gene ontology (GO) and KEGG enrichment

Official gene IDs for transcripts that were differentially modulated following HPAI H5N1 infection with or without apocynin treatment were compiled into three target lists. Statistically significant differentially expressed transcripts were defined as having ≥ 2-fold change with a Benjamini-Hochberg adjusted *P* value < 0.01. A background list of genes was compiled by retrieving all gene IDs identified from the present HiSeq analysis with FPKM > 1. Biological process GO enrichment was performed using GOrilla comparing unranked background and target lists (58) and redundant GO terms were removed using REVIGO (59). The target lists were also subjected to KEGG pathway analysis using a basic KEGG pathway mapper (60).

### Quantitative real-time reverse transcriptase polymerase chain reaction (qRT-PCR)

mRNA concentrations of genes of interest were assessed and analyzed using qRT-PCR as previously described (5). Briefly, after total RNA extraction from influenza-infected cells, cDNA was prepared using SuperScript™ III First-Strand Synthesis SuperMix (Invitrogen). Gene expression levels of various cytokines were assessed using TaqMan Gene Expression Assays (Applied Biosystems) with commercial TaqMan primers and probes, with the exception of influenza Matrix (M) gene (forward primer 5’-CTTCTAACCGAGGTCGAAACGTA-3’; reverse primer 5’-GGTGACAGGATTGGTCTTGTCTTTA-3’; probe 5’-FAM-TCAGGCCCCCTCAAAGCCGAG-NFQ-3’) (61). Specific primers (Table S5) were designed to estimate the expression of *CEACAM1* variants in ATII and A549 cells using iTaq Universal SYBR Green Supermix (BioRad) according to manufacturer’s instruction. The absence of nonspecific amplification was confirmed by agarose gel electrophoresis of qRT-PCR products (15 μL). Gene expression was normalized to β-actin mRNA using the 2^−ΔΔCT^ method where expression levels were determined relative to uninfected cell controls. qRT-PCR primers and primer pairs used to identify each *CEACAM1* variant are listed in Table S5. All assays were performed in duplicate using an Applied Biosystems^®^ StepOnePlus™ Real-Time PCR System.

### Western blot analysis

Protein expression of CEACAM1 was determined using Western blot analysis as previously described (5). Protein concentrations in cell lysates were determined using EZQ^®^ Protein Quantitation Kit (Molecular Probes™, Invitrogen). Equal amount of proteins was loaded and resolved by SDS/PAGE and transferred to PVDF membranes (BioRad). Membranes were probed with rabbit anti-CEACAM1 monoclonal antibody (EPR4049 (ab108397), Abcam). Proteins were visualized using Pierce enhanced chemiluminescence (ECL) Plus Western Blotting Substrate (Thermo Scientific). Protein band intensity was quantified using Fiji software (version 1.49J10) (62), normalized against β-actin as a loading control (rabbit monoclonal, HRP-conjugated, Cell Signaling), and expressed as fold changes compared to controls.

### Knockdown of endogenous CEACAM1

ATII and A549 cells were grown to 80% confluency in 6-well plates then transfected with small interfering RNA (siRNA) targeting the human *CEACAM1* gene (siCEACAM1; s1976, Silencer^®^ Select Pre-designed siRNA, Ambion^®^) or siRNA control (siNeg; Silencer^®^ Select Negative Control No. 1 siRNA, Ambion^®^) using Lipofetamine 3000 (ThermoFisher Scientific) according to manufacturer’s instructions. Transfection and silencing efficiency were evaluated after 48 hours by Western blot analysis of CEACAM1 protein expression and by qRT-PCR analysis of *CEACAM1* variants. In parallel experiments, virus replication and cytokine/chemokine production were analyzed in siCEACAM1-or siNeg-transfected cells infected with H5N1 virus (MOI = 0.01) at 24 hpi.

### Statistical analysis

Differences between two experimental groups were evaluated by using Student’s unpaired, two-tailed *t* test. Differences in fold-changes of mRNA expression (qRT-PCR) between three experimental groups were evaluated using one-way analysis of variance (ANOVA) followed by a Bonferroni multiple-comparison test. Differences were considered significant with a *p* value of < 0.05. The data are shown as means ± standard error of the mean (SEM) from three or four individual experiments. Statistical analysis was performed using GraphPad Prism for Windows (v5.02).

## ACKNOWLEDGMENTS

This study was supported by funding from The Molecular and Medical Research Strategic Research Centre (MMR SRC), Deakin University and The Geelong Centre for Emerging Infectious Diseases (GCEID) (to S.Y.), and by an Alfred Deakin Postdoctoral Research Fellowship (to S.Y.).

The authors wish to thank AAHL CSIRO and the University of Melbourne for providing viruses. We thank Katie-Lee Alexander from Barwon Health Tissue Bank, and Samantha Arandelovic from St John of God Pathology, Geelong, for the coordination of human lung tissue collection and sampling.

The authors declare no conflict of interest.

